# Giantin knockout models reveal a feedback loop between Golgi function and glycosyltransferase expression

**DOI:** 10.1101/123547

**Authors:** Nicola L. Stevenson, Dylan J. M. Bergen, Roderick E.H. Skinner, Erika Kague, Elizabeth Martin-Silverstone, Kate A. Robson Brown, Chrissy L. Hammond, David J. Stephens

## Abstract

The Golgi is the cellular hub for complex glycosylation, controlling accurate processing of complex proteoglycans, receptors, ligands, and glycolipids. Its structure and organisation is dependent on golgins, which tether cisternal membranes and incoming transport vesicles. Here we show that knockout of the largest golgin, giantin, leads to substantial changes in gene expression despite only limited effects on Golgi structure. Notably, 22 Golgi-resident glycosyltransferases, but not glycan processing enzymes or the ER glycosylation machinery, are differentially expressed following giantin ablation. This includes near-complete loss-of-function of GALNT3 in both mammalian cell and zebrafish models. Giantin knockout zebrafish exhibit hyperostosis and ectopic calcium deposits, recapitulating phenotypes of hyperphosphatemic familial tumoral calcinosis, a disease caused by mutations in GALNT3. These data reveal a new feature of Golgi homeostasis, the ability to regulate glycosyltransferase expression to generate a functional proteoglycome.

**Summary statement:** Knockout of giantin in a genome-engineered cell line and zebrafish models reveals the capacity of the Golgi to control its own biochemistry through changes in gene expression.

## Introduction

Golgins are coiled-coil domain proteins that project out from the surface of the Golgi apparatus into the cytosol (Gillingham and Munro, 2016). They maintain Golgi organisation and selectively tether incoming transport vesicles seeking to fuse with Golgi cisternae. The largest golgin family member is giantin, whose N-terminal cytosolic domain has a predicted molecular mass of 370kDa (Linstedt and Hauri, 1993). Giantin is one of only three golgins to have a C-terminal transmembrane domain, directly anchoring it within cis-and medial-Golgi membranes.

The functional role of giantin is poorly defined. Early *in vitro* studies suggest that giantin resides in COPI vesicles; transport carriers mediating intra-Golgi and retrograde Golgi-to-endoplasmic reticulum (ER) transport (Sönnichsen et al., 1998). Here, giantin is reported to recruit p115, which binds simultaneously to GM130 on cis-Golgi membranes to mediate tethering. Giantin-p115 interactions may also facilitate GM130-independent retrograde transport (Alvarez et al., 2001). In addition to p115, giantin has been shown to interact with GCP60 (Sohda et al., 2001), Rab1, and Rab6 (Rosing et al., 2007). Rab1 and Rab6 localise to ER-Golgi- and retrograde transport vesicles respectively and thus their interaction with Golgi-resident giantin could similarly promote vesicle capture. Furthermore, giantin is also implicated in lateral Golgi tethering (Koreishi et al., 2013) and ciliogenesis (Asante et al., 2013; Bergen et al., 2017).

Rodent models carrying loss-of function alleles of giantin vary in phenotype. Homozygous knockout (KO) rats, possessing a null mutation in the *Golgb1* gene encoding giantin, develop late embryonic lethal osteochondrodysplasia (Katayama et al., 2011). Embryonic phenotypes include systemic oedema, cleft palate, craniofacial defects, and shortened long bones which are largely attributed to defects in chondrogenesis. Interestingly, chondrocytes from homozygous animals have expanded ER and Golgi membranes whilst cartilage growth plates contain less extracellular matrix (ECM), indicative of secretory pathway defects (Katayama et al., 2011). Mouse giantin KO models have less complex developmental disorders, the most predominant phenotype being cleft palate (Lan et al., 2016) and short stature (McGee et al., 2017). These animals also have ECM abnormalities associated with glycosylation defects but Golgi structure is normal (Lan et al., 2016). Work from our lab has also now characterised giantin function in zebrafish (Bergen et al., 2017). In contrast to rodent models, homozygous giantin KO zebrafish do not show any gross morphological changes during development, can reach adulthood and show only a minor growth delay. They do however show defects in cilia length consistent with our previous work *in vitro* (Asante et al., 2013). We have also defined defects in procollagen secretion following RNAi of giantin expression in cultured cells (McCaughey et al., 2016). Thus, defects in ECM assembly could underpin some of the developmental defects seen in giantin knockout model organisms.

There are two major pathways of protein glycosylation, *N*- and *O*-glycosylation initiated in the ER and Golgi respectively. Most oligosaccharides are then subject to modification and extension by Golgi-resident type II transmembrane glycosyltransferases, the importance of which is underscored by the clear link between Golgi dysfunction and congenital disorders of glycosylation (Freeze and Ng, 2011). Mucin-type O-glycosylation is the most prevalent form of glycosylation on cell surface and secreted proteins. It is initiated by Golgi-resident polypeptide *N*-acetylgalactosaminyltransferases (GALNTs) that catalyse the addition of *N*-acetylgalactosamine to serine or threonine residues on target substrates (forming the Tn antigen, (Bennett et al., 2012)). There are twenty GALNT proteins in humans with distinct but overlapping substrate specificities and spatio-temporal expression patterns (Bard and Chia, 2016; Schjoldager et al., 2015). Such redundancy means mutations in GALNT genes generally produce very mild phenotypes, although several genome-wide association studies have linked GALNTs with diverse pathologies such as Alzheimer’s disease (Beecham et al., 2014) and obesity (Ng et al., 2012). Moreover, bi-allelic loss-of function mutations in GALNT3 have been directly linked to the human disease hyperphosphatemic familial tumoral calcinosis (HFTC, (Ichikawa et al., 2007; Kato et al., 2006; Topaz et al., 2004). In such cases, complete loss of GALNT3 function results in a failure to *O*-glycosylate FGF23, leading to its inactivation and the subsequent development of hyperostosis and ectopic calcium deposits in skin and subcutaneous tissues.

In the absence of a clearly defined role for giantin at the Golgi, we sought to study its function in an engineered KO cell line. In this system, as well as a zebrafish model, we show for the first time that loss of giantin results in changes in the expression Golgi-resident glycosyltransferases, defining a new role for giantin in quality control of Golgi function through transcriptional control.

## Results

### Generation of a giantin KO cell line

We generated a KO cell line for *GOLGB1* (giantin) using genome editing. A GFP-fusion of the double nickase mutant of Cas9 (Cas9^D10A^-GFP) was co-transfected into human non-transformed telomerase immortalized retinal pigment epithelial (hTERT-RPE-1) cells with paired guide RNAs targeting exon 7 of the *GOLGB1* gene. GFP-positive cells were then sorted by fluorescence activated cell sorting, screened for loss of giantin by immunofluorescence, and sequenced at the target site. Using this approach, one clone was identified with an indel frameshift mutation in both alleles, leading to a frameshift and premature stop codon in exon 4 (full annotation: R195fsX204-R195P-A196del, Figure 1A). Downstream of the mutation an in-frame translational start site was also noted with the potential to permit expression of a truncated protein. To exclude this possibility, we probed the mutant cells for giantin expression using three different antibodies raised against the full length, C-, and N-termini of the protein. No protein was detected by immunoblot or immunofluorescence using these antibodies (Figure 1B-D).

**Figure 1:**
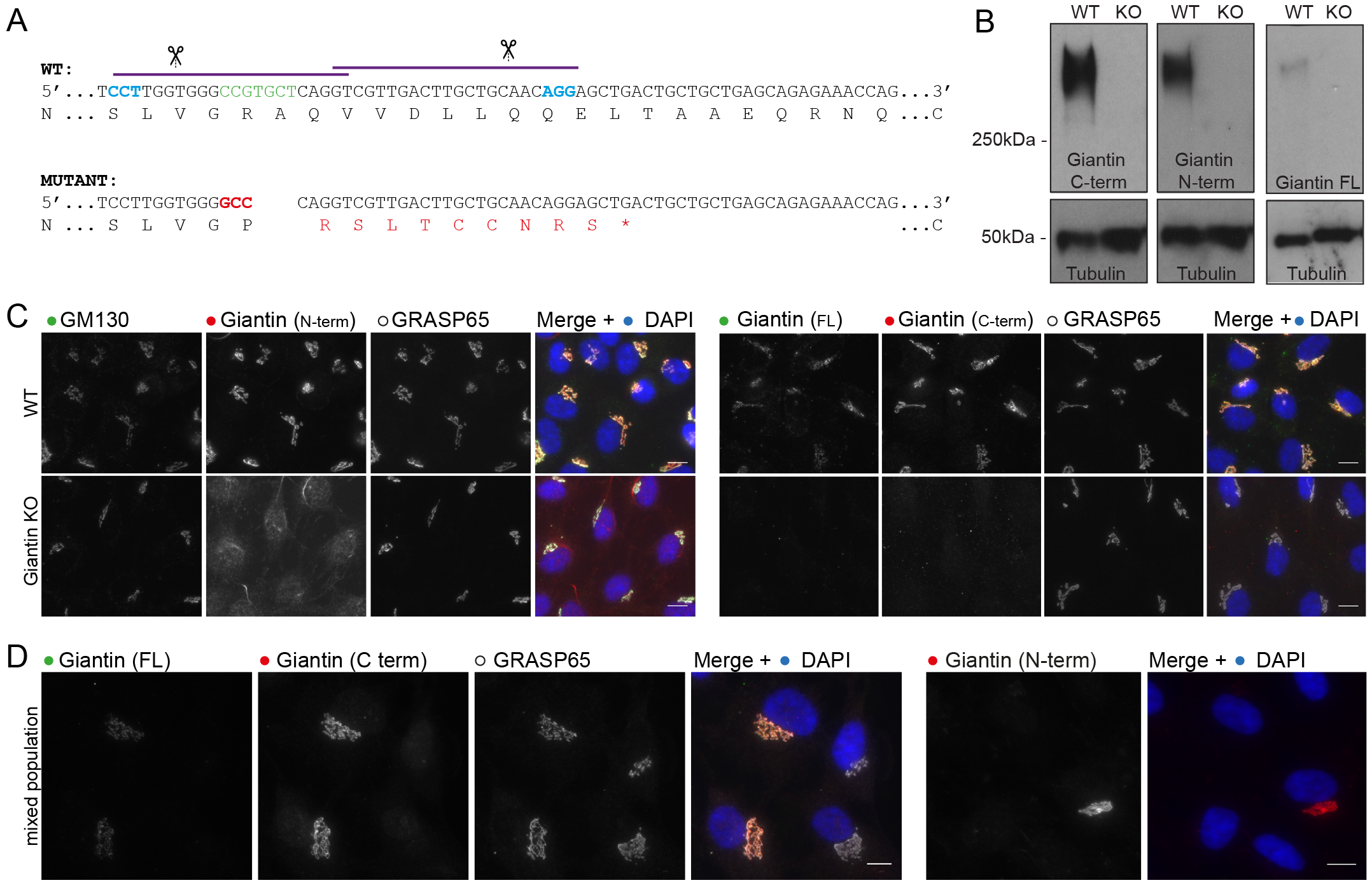
Generation of a giantin KO cell line. A. Genomic sequence for CRISPR-Cas9 target site in WT and engineered KO RPE-1 cell line. Purple lines and scissors depict gRNA binding and cut sites. Blue nucleotides show the CRISPR PAM site. Green and red nucleotides are those deleted and inserted in the KO mutation respectively. Amino acid translation shown underneath; asterisk indicates a premature stop codon. B. Western blot analysis and C. immunofluorescence staining of giantin using three different antibodies raised against the C-terminus (C-term), N-terminus (N-term) and full length (FL) protein. All immunoreactivity is lost in the KO cells. D. WT and KO cell mixed population stained for giantin and other Golgi markers for direct comparison. Images are maximum projections. Scale bars 10μm.

### Loss of giantin does not lead to gross defects in Golgi morphology or trafficking

As giantin resides at the Golgi apparatus, we began characterising the KO cell line by examining Golgi morphology. KO cells were immuno-labelled for Golgi markers and the size and number of Golgi elements were quantified. No significant change in Golgi structure was detected (Figure 2A-C). The relative distribution of *cis*- and *trans*-Golgi markers was also maintained, suggesting organelle polarity was unperturbed (Figure 2D). Similarly, the general organisation of the early secretory pathway was normal (Figure 2E, showing labelling for ER exit sites and ER-Golgi intermediate compartment). We therefore decided to study Golgi morphology in greater detail by electron microscopy (EM). At this resolution, Golgi stacks had comparable numbers of cisternae in WT and KO cells and cisternae were of equivalent length with no sign of dilation (Figure 2F-H). Overall these results suggest Golgi structure was not grossly disrupted following loss of giantin.

**Figure 2:**
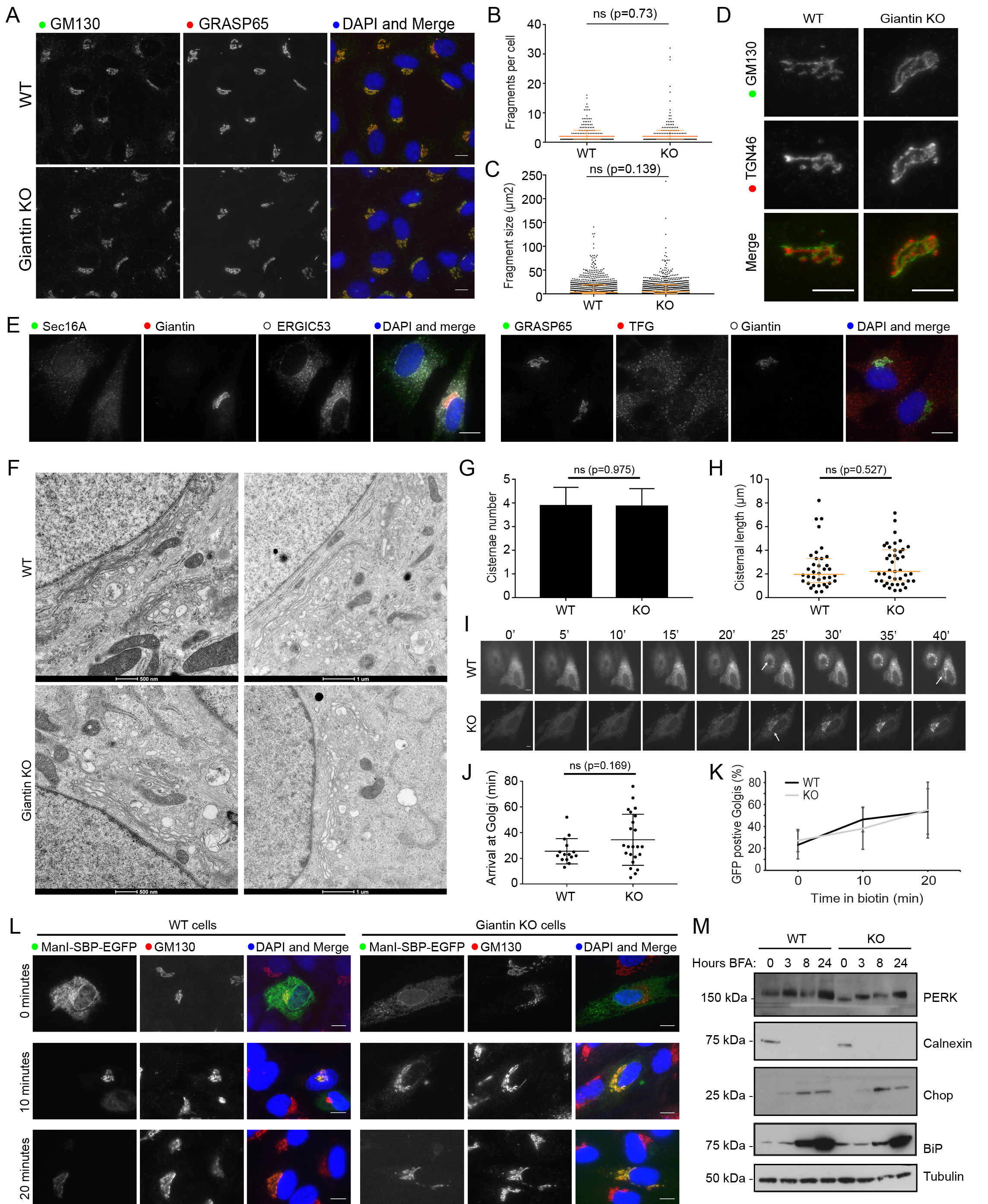
Loss of giantin has no effect on Golgi structure or trafficking. A. Representative images of WT and KO cells immuno-labelled for two *cis*-Golgi markers. The number of GM130 positive elements per cell (B) and their area (C) was found to be equivalent in WT and KO cells (n=3; 387 WT and 320 KO cells quantified; orange bars indicate median and interquartile range; statistics Mann-Whitney; fragments smaller than 0.5μm^2^ excluded). D. Co-labelling of cells with *cis*- (GM130) and *trans*- (TGN46) Golgi markers shows Golgi polarity is maintained in KO cells. E. Representative images of WT and KO cells immuno-labelled for early secretory pathway markers as indicated. A-E. Images shown are maximum projections. Scale bar 10μm. F. Transmission electron micrographs of Golgi elements in WT and KO cells. The number of cisternae per stack (G) and length of Golgi cisternae (H) was quantified from experiments represented in (F) (n=3; total 30 cells per cell line; orange bars indicate median and interquartile range; statistics Mann-Whitney). I-L. WT and KO cells expressing Str-Kdel/ManII-SBP-EGFP were treated with biotin and imaged live (I-J) or fixed at 0, 10, and 20 minutes post-biotin addition and immuno-labelled for GM130 (K-L). I. Single plane images taken from representative movies at 5-minute intervals. See supplementary movies. Scale bar 10μm. Arrows show arrival of reporter at Golgi. J. Quantification of the time at which fluorescence appears in the Golgi apparatus in movies represented in (I) (n=3; 15 WT cells and 23 KO cells quantified; bars show median and interquartile range; statistics Mann-Whitney). K. Quantification of the number of GFP-positive Golgi at each timepoint in fixed cells (n=3; 378 WT and 310 KO cells quantified; mean and standard deviation shown). L. Representative single plane images of fixed cells at each timepoint. M. Western blot analyses of ER stress markers in lysates taken from WT and KO cells following treatment with BFA for the indicated time.

Many golgins have been shown to act as tethers for transport vesicles but such a function has not yet been defined for giantin (Wong and Munro, 2014). To test whether giantin is involved in trafficking, we used the Retention Using Selective Hooks (RUSH) system (Boncompain et al., 2012) to monitor ER-to-Golgi transport. In this assay a fluorescently-labelled Golgi-resident protein (the reporter, here EGFP-tagged mannosidase II) is fused to streptavidin binding protein (SBP) and co-expressed with an ER-resident protein fused to streptavidin (the hook, here tagged with a KDEL motif). When both engineered fusion proteins are present, the SBP on the reporter binds to the streptavidin on the hook and is retained in the ER. Reporter release is then induced by the addition of biotin, which outcompetes the SBP for streptavidin binding. Time-lapse imaging of biotin treated KO cells expressing this RUSH construct (Supplementary Movies S1 and S2) showed a slight delay and greater variability in anterograde mannosidase-II trafficking relative to WT, however this difference was not statistically significant (Figure 2I-J). In order to analyse a greater number of cells, we repeated this experiment in fixed cells at 0, 10, and 20 minutes post-biotin addition and quantified cargo delivery at each time point. Again, giantin KO cells showed no significant delay in anterograde transport compared to WT cells (Figure 2K-L). This approach also allowed us to confirm that we were observing ER-Golgi transport as we could co-label the Golgi (Figure 2L).

Perturbations in anterograde trafficking can result in ER stress and activation of the unfolded protein response (UPR) as secretory cargo accumulates in this compartment (Brodsky, 2012). We found that expression of UPR markers including PERK, calnexin and CHOP were unchanged in giantin KO cells compared to controls (Figure 2M), suggesting no activation of the UPR in giantin KO cells.

### GM130 localisation is altered in giantin KO cells following Golgi fragmentation

During mitosis, the Golgi must disassemble and reassemble. As we could not detect any gross defects in Golgi structure in giantin KO cells at steady state, we analysed Golgi dynamics by chemically inducing its disassembly. First, we treated cells with nocodazole, which disassembles microtubules and thus causes Golgi ribbons to fragment into polarised mini-stacks (Thyberg and Moskalewski, 1985). Under these conditions, the dynamics of disassembly and reassembly were found to be equivalent in both cell lines (Figure 3A), with fragmentation of the TGN preceding that of the cis-Golgi as reported previously (Yang and Storrie, 1998). Likewise, Golgi disassembly following brefeldin A treatment (which inhibits the Arf-guanine nucleotide exchange factor, GBF1) was comparable in WT and KO cells (Figure S1). We also failed to find any defects in cell cycle progression using propidium iodide labelling and flow cytometry (data not shown).

**Figure 3:**
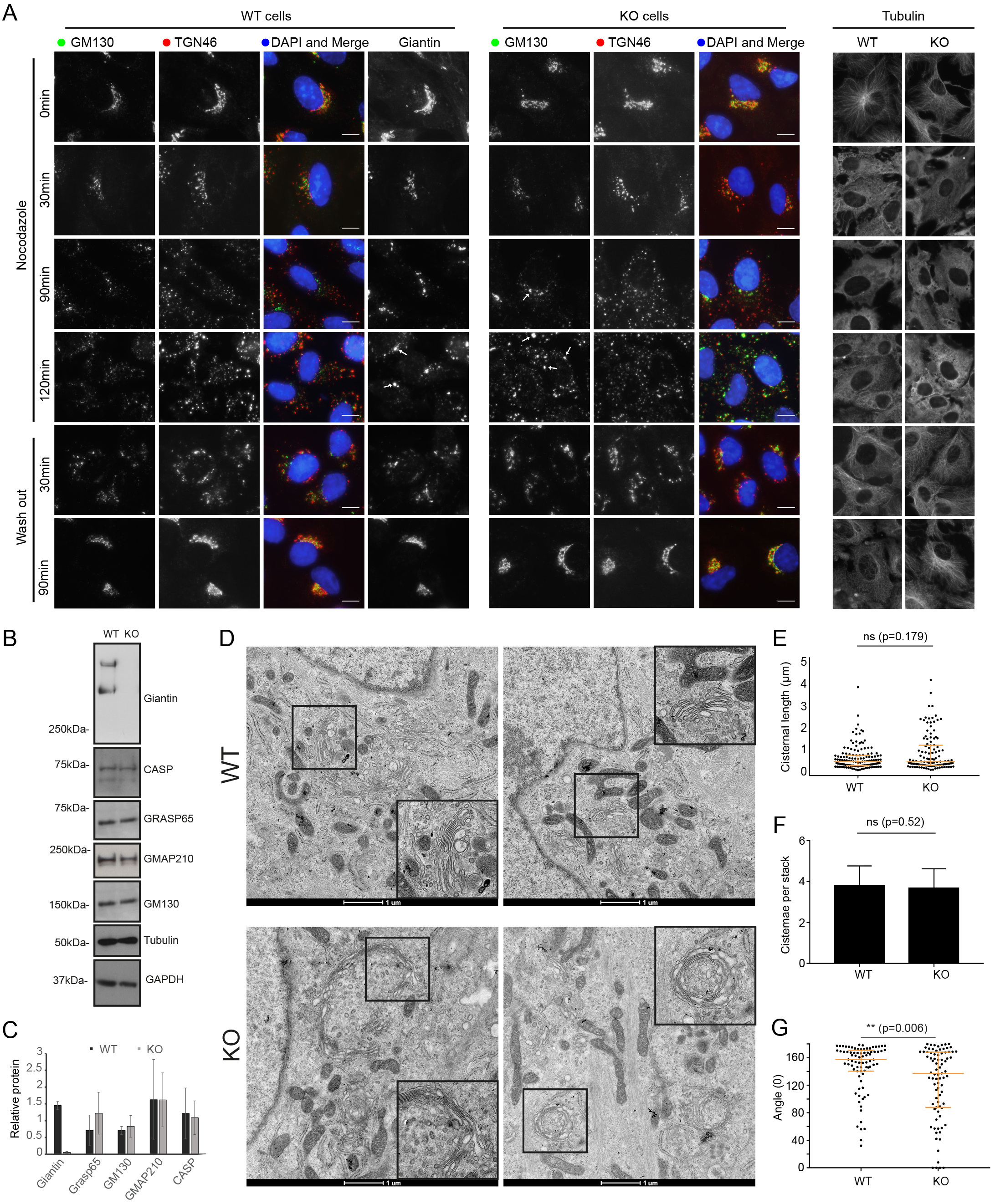
Giantin loss leads to mild changes in Golgi mini-stack structure. A. Representative maximum projection images of WT and giantin KO cells incubated with 5μm nocodazole as indicated and immuno-labelled for *cis*- (GM130) and *trans*- (TGN46) Golgi markers or tubulin. In wash out panels, cells were incubated with nocodazole for 3 hours then washed and incubated in growth medium for time indicated. Scale bars 10μm. B. Western blot analysis of golgin expression in WT and KO cells. C. Quantification of blots represented in (B) (n=3, mean and standard deviation shown). D. Transmission micrographs of WT and KO cells incubated with 5μm nocodazole for 90 minutes. Inserts show zoom of region denoted by black squares. E-G. Quantification of experiments represented in D showing (E) cisternal length, (F) number of cisternae per stack and (G) the angle between lines drawn from each lateral rim of the stack to the centre (n=3; 27 WT and 21 KO cells quantified; E and G show median and interquartile range, F mean and standard deviation; statistics Mann-Whitney).

During these Golgi disruption experiments, we noticed a difference in GM130 labelling of WT and KO cells. Following nocodazole treatment, giantin reportedly persists on the original fragmenting membranes (the ‘old Golgi’) rather than cycling through the ER onto immature peripheral mini-stacks (Fourriere et al., 2016; Nizak et al., 2003). This is apparent here in WT cells, which show an enrichment of giantin on larger, juxtanuclear structures over more peripheral elements (Figure 3A). In KO cells however, these larger Golgi elements are enriched with GM130. This enrichment is not due to upregulation of GM130 expression, as protein levels are equivalent in WT and KO cells (Figure 3B-C) suggesting instead that GM130 has either redistributed between Golgi membranes, perhaps to compensate for giantin, or is labelling larger structures not present in WT cells. To distinguish between these possibilities, we examined cells treated with nocodazole for 90 minutes by EM. As expected larger, perinuclear ‘old Golgi’ structures could be detected in both WT and KO cells, as well as peripheral mini-stacks (Figure 3D). The size distribution of these structures was equivalent in both cell lines (Figure 3E-F) indicating the larger GM130-labelled elements reflect a redistribution of the protein.

### Giantin negative Golgi ‘mini-stacks’ show a tendency to circularise

Surprisingly, EM of nocodazole-treated KO cells showed Golgi elements that had apparently circularised (Figure 3D, insets). These were absent in WT and untreated KO cells, except for one case of the latter. To quantify curvature of fragmented Golgi elements we calculated the angle between two lines drawn from each Golgi rim to the centre of the stack; circularised Golgi structures were assigned an angle of 0° and linear stacks 180°. This analysis showed a significant overall trend towards horseshoe-shaped and circular stacks in the KO cells compared to the WT (Figure 3G). Giantin-deficient Golgi stacks therefore exhibit structural abnormalities with low frequency (5% of structures/at least one present in 14% of cells) once fragmented.

### Glycosylation enzyme expression patterns are altered in giantin KO cells

Giantin is an evolutionarily conserved gene essential for viability in rodents (Katayama et al., 2011; Lan et al., 2016), yet phenotypes in our KO cell line and indeed in KO zebrafish (Bergen et al., 2017) are mild. We therefore considered whether the cells had undergone adaptation, as has been reported for other KO systems (Rossi et al., 2015). Having established that the expression of other golgin family members was normal (Figure 3B-C), we performed RNAseq of WT and KO cells to compare gene expression patterns in an unbiased manner. Pairwise analysis of triplicate samples identified a total of 1519 genes showing a greater than 2-fold change in expression in KO cells. Of those, 807 genes exhibited a greater than 3-fold change in expression in KO cells (Supplementary Table S1). Gene ontology analysis showed that the major classes of genes that were differentially expressed encoded highly glycosylated proteins, extracellular matrix components, and adhesion proteins. Twenty-four glycosyltransferases were differentially expressed between the two cell lines. These include a pseudogene (DPY19L2P2), an ER-resident glycosyltransferase (UGT8) and twenty-two type II Golgi-resident transmembrane enzymes (Table 1). Some of these were among the most highly downregulated genes overall. Notably, other ER-localised core glycosyltransferases, glycan processing and modifying enzymes, and the cytosolic glycosylation machinery were unchanged following KO of giantin.

**Table 1:** Glycosylation enzymes differentially expressed between WT and giantin KO RPE-1 cells. Values shown are Fragments Per Kilobase of transcript per Million mapped reads (FPKM), the log_2_-fold change between these and the uncorrected p- and q-values (q being the false discovery rate, also known as (FDR)-adjusted p-value). All values were found significant (where p is greater than the FDR after Benjamini-Hochberg correction for multiple-testing). Pathway annotation and steady-state localisation was done manually based on gene ontology and published literature. Genes highlighted in red are downregulated and green are upregulated.

| Gene | WT FPKM | KO FPKM | log2 (fold change) | p_value | q_value | Pathway | Organelle |
| --- | --- | --- | --- | --- | --- | --- | --- |
| GALNT5 | 0.0293487 | 0.693322 | 4.56216 | 5.00E-05 | 0.000214 | N-Glycosylation | Golgi |
| ST6GALNAC3 | 0.302522 | 2.43643 | 3.00966 | 5.00E-05 | 0.000214 | O-Glycosylation | Golgi |
| EXTL1 | 0.417023 | 1.00605 | 1.2705 | 5.00E-05 | 0.000214 | N-Glycosylation | Golgi |
| CSPG5 | 0.351491 | 0.173152 | -1.02145 | 0.0001 | 0.000408 | O-Glycosylation | Golgi |
| GAL3ST3 | 0.252928 | 0.123266 | -1.03695 | 0.00015 | 0.000591 | Both N- and O-Glycosylation | Golgi |
| GALNT1 | 153.498 | 71.0283 | -1.11175 | 5.00E-05 | 0.000214 | O-Glycosylation | Golgi |
| DPY19L2P2 | 1.60655 | 0.718415 | -1.16107 | 5.00E-05 | 0.000214 | C-Glycosylation | Pseudogene |
| ST6GALNAC2 | 0.595085 | 0.24238 | -1.29582 | 5.00E-05 | 0.000214 | N-Glycosylation | Golgi |
| MGAT5B | 6.70667 | 2.5532 | -1.39329 | 5.00E-05 | 0.000214 | N-Glycosylation | Golgi |
| GALNT16 | 9.52671 | 3.35166 | -1.5071 | 5.00E-05 | 0.000214 | O-Glycosylation | Golgi |
| B4GALNT4 | 9.7297 | 3.34711 | -1.53948 | 5.00E-05 | 0.000214 | O-Glycosylation | Golgi |
| B3GNT5 | 1.77188 | 0.581017 | -1.60863 | 5.00E-05 | 0.000214 | O-Glycosylation | Golgi |
| A4GALT | 6.16651 | 1.7832 | -1.78999 | 5.00E-05 | 0.000214 | O-Glycosylation | Golgi |
| HS3ST1 | 1.92483 | 0.54196 | -1.82847 | 5.00E-05 | 0.000214 | O-Glycosylation | Golgi |
| LFNG | 2.58988 | 0.653085 | -1.98754 | 5.00E-05 | 0.000214 | Both N- and O-Glycosylation | Golgi |
| CHST11 | 9.46678 | 1.97911 | -2.25802 | 5.00E-05 | 0.000214 | O-Glycosylation | Golgi |
| CHSY3 | 0.8767 | 0.161329 | -2.44208 | 5.00E-05 | 0.000214 | O-Glycosylation | Golgi |
| GALNT12 | 1.42884 | 0.199249 | -2.84221 | 5.00E-05 | 0.000214 | O-Glycosylation | Golgi |
| GBGT1 | 2.71611 | 0.233266 | -3.54149 | 5.00E-05 | 0.000214 | Glycolipid Glycosylation | Golgi |
| FUT4 | 0.904089 | 0.0770361 | -3.55286 | 5.00E-05 | 0.000214 | N-Glycosylation | Golgi |
| UGT8 | 4.82991 | 0.272601 | -4.14714 | 5.00E-05 | 0.000214 | Glycolipid Glycosylation | ER |
| GALNT3 | 8.35694 | 0.33148 | -4.65598 | 5.00E-05 | 0.000214 | O-Glycosylation | Golgi |
| ST6GAL2 | 0.699234 | 0.0208263 | -5.0693 | 5.00E-05 | 0.000214 | N-Glycosylation | Golgi |
| ST8SIA4 | 0.341234 | 0 | - | 5.00E-05 | 0.000214 | N-Glycosylation | Golgi |

To determine the impact of altered glycosyltransferase expression in the KO cells, we looked at global glycosylation patterns using biotinylated lectins to label fixed cells. RCA_120_ labelling of β-D-galactosyl residues was more bundled in KO cells, but otherwise there were no gross changes in glycan abundance or localisation (Figure S2). We also probed cell lysates with lectins by blotting and found only minor changes in glycosylation patterns, namely loss of a 25 kDa band when labelling with either ConA or HABP which recognise α-D-mannosyl and α-D-glucosyl residues and hyaluronic acid respectively (Figure S2). Glycosylation patterns are therefore largely normal, but with some identifiable changes.

### GALNT3 expression is substantially reduced in giantin KO cells

To validate the findings of the RNAseq analysis, we first performed an immunoblot for one of the more highly downregulated glycosyltransferases, GALNT3. This confirmed that protein expression was almost completely abolished in 5 biological replicates (Figure 4A). Immunolabelling of fixed cells further demonstrated a near-complete loss of GALNT3 expression in giantin KO cells (Figure 4B). Additionally, investigation into the most highly upregulated gene overall, RCAN2, similarly corroborated the RNAseq results at the protein level (Stevenson et al., 2017).

**Figure 4:**
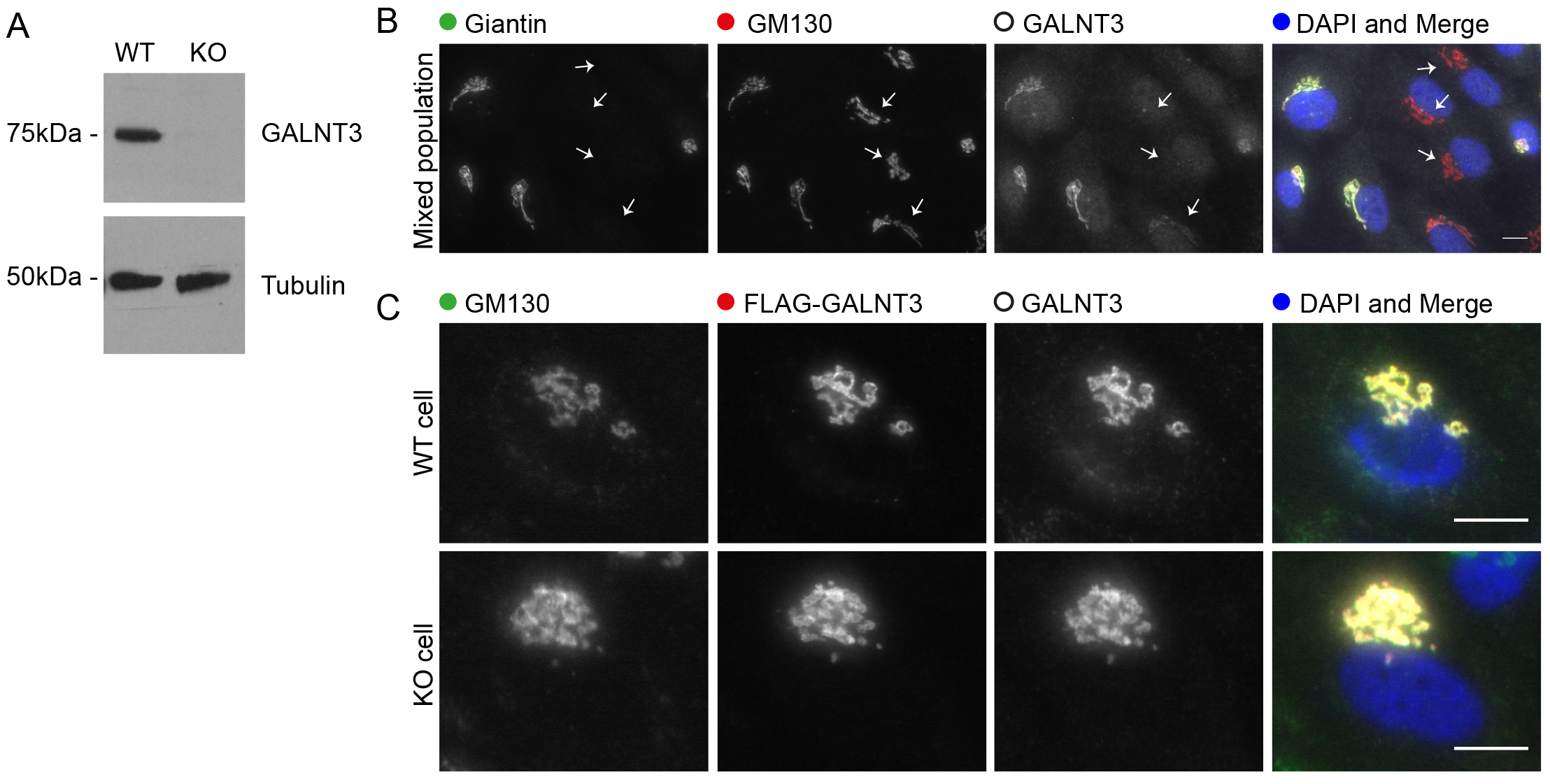
GALNT3 expression is lost in giantin KO cells. A. Western b 551 lot validating down-regulation of GALNT3 in KO cells (n=5 biological replicates). B. Maximum projection images of mixed populations of WT and KO cells immuno-labelled for giantin, GM130 and GALNT3. Arrows highlight giantin KO cells. C. Representative projections of WT and KO cells expressing FLAG-tagged GALNT3 fixed and stained as indicated. All scale bars 10μm.

*GALNT3* is mutated in the human disease HFTC (Topaz et al., 2004) and so we decided to focus our studies on this gene. We hypothesised that downregulation of GALNT3 could have occurred in response to aberrant trafficking following the loss of giantin function; such mistargeting could result in degradation coupled with a feedback mechanism to downregulate expression. We tested this directly by expressing FLAG-tagged GALNT3 in WT and KO cells. Immunofluorescence labelling showed FLAG-GALNT3 is efficiently targeted to the Golgi in both cell lines (Figure 4C). GALNT3 localisation is thus independent of giantin function and not the cause of its down-regulation. We next decided to test whether GALNT3 down-regulation was reversible by reintroducing epitope-tagged giantin into KO cells. Giantin KO cells expressing flag-giantin for up to 2 weeks failed to show any recovery of GALNT3 protein expression (data not shown).

### Giantin KO zebrafish phenotypes are consistent with tumoral calcinosis

We next sought to explore the role of giantin in regulating glycosyltransferase expression *in vivo*, using two recently characterised *golgb1* KO zebrafish lines (Bergen et al., 2017). The first of these, derived by ENU mutagenesis, carries a point mutation (C>T) in exon 14 leading to generation of a premature stop codon at glutamine-2948 (denoted *golgb1^Q2948X/Q2948X^*). The second allele was generated by TALEN site-directed mutagenesis, introducing an 8bp insertion at exon 14. This results in a frameshift at position 3029 leading to a premature stop codon at position 3078 (E3027fsX3078-T3028_A3029del, denoted *golgb1^X3078/X3078^*). Both mutations lead to loss of the transmembrane domain and therefore are expected to be loss-of-function mutations. These fish do not display any gross developmental defects, but did have a mild developmental delay and an increase in cilia length (Bergen et al., 2017).

Given the skeletal defects seen in human patients lacking GALNT3 function and in giantin KO rodents, we performed quantitative PCR of mixed bone and cartilage tissues from both mutant fish lines at 60 days post fertilisation (dpf). In each case, we observed a significant reduction of *galnt3* expression (Figure 5A) with one KO individual from each line possessing almost undetectable levels of transcript. Since the giantin KO fish reach adulthood, and given the causative link between loss of GALNT3 and HFTC in humans, we next examined WT and mutant skeletal structures by micro-computed tomography (micro CT) in both mutant lines; 3 mutants and siblings of the Q2948X line were scanned at 8 months post fertilisation and 4 mutants and siblings of the X3078 line were scanned at 10 months post fertilisation. Both mutant lines showed relatively normal skeletal patterning consistent with the published phenotype (Figure 5B-I and (Bergen et al., 2017)). However, both lines exhibited ectopic mineralisation of soft tissues. All 4 mutants from the X3078 line showed ectopic mineralisation of the intervertebral discs, leading to reduction in vertebral spacing and vertebral fusions (Figure 5 B, C). Furthermore, 2 out of 3 *golgb1*^*Q2948X/Q2948X*^ adults showed ectopic calcium-like deposits in the soft tissues of the thoracic cavity near bone elements (Figure 5D G and Supplementary Movies S3 and S4), while the 3^rd^ showed deposits within multiple vertebrae (Figure 5I, and Supplementary Movies S5 and S6). In addition to aberrant mineralisation, HFTC is also associated with hyperostosis. In addition to the fused vertebrae in both mutant lines we also observed craniofacial alterations such that the lower jaw was longer and narrower than in their wild type siblings (Figure S3), consistent with altered bone deposition (Figure S3). Total bone mineral density was not significantly different between the mutants and siblings. However, when readings were taken across large volumes in the jaw, skull or vertebrae, we observed higher variability in the mutants than in the wild type fish. This is also consistent with alterations to skeletal homeostasis (Figure S3).

**Figure 5:**
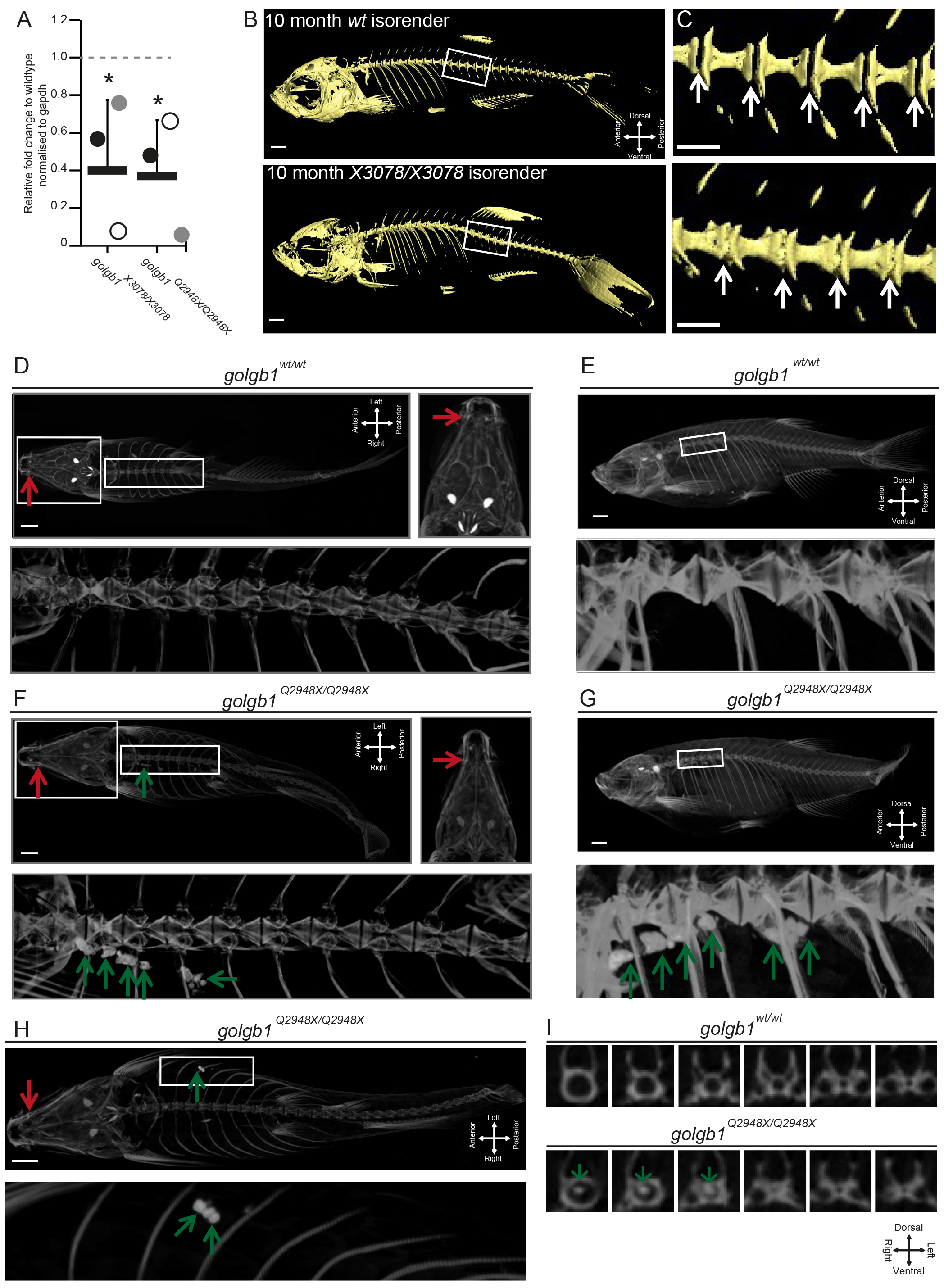
Giantin KO zebrafish have reduced galnt3 expression and exhibit HFTC-like phenotypes. A. Real-time qPCR pairwise analysis of *galnt3* expression at 60-63 dpf in two *golgbl* mutant zebrafish lines normalised to *gapdh* mRNA levels as housekeeping gene. Bars show mean expression for each mutant line (n=3 per genotype group) relative to WT siblings (WT expression 1A.U. depicted by dashed line). Each circle represents one individual (P value: *=<0.05, mean with standard deviation). B. Lateral views of micro CT scans of 10-month WT and golgb1^X3078/X3078^ homozygous mutants, presented as isosurface renders. Boxed regions show enlarged regions in C. C Enlarged regions of the spine; white arrows demarcate intervertebral discs (IVDs), which in wild type are not mineralised but in the mutant ectopic mineralisation is seen manifesting as vertebral fusions. D. Ventral (with high resolution inset) and E. lateral view micro CT images showing craniofacial and spinal elements of a representative WT sibling (Q2948X line, n=3 females). F-I. Three golgb1^Q2948X/Q2948X^ female individuals showing ectopic calcium deposits in soft tissues (F. ventral view with high resolution inset and G. lateral view of individual 1, and H. ventral view of individual 2) and in spinal column (I. digital axial z-slices of individual 3). D-I. Red arrows indicate mandible joint and green arrows ectopic deposits. Line Q2948X were imaged at 8 months post fertilisation. (A) Unpaired t-test was used as data were normally distributed. Scale bar 100μm.

## Discussion

The data presented here demonstrate for the first time that the Golgi apparatus has the capacity to control its own enzymatic composition through changes in transcription. Specifically, we show that the enzymatic content of the Golgi is altered at the level of transcription in response to loss of giantin function. The ER and cytosolic glycosylation machineries, as well as Golgi-localised glycosidases, are unaffected. This process is conserved between mammalian cells and zebrafish models as GALNT3 mRNA is reduced in both giantin KO systems. Furthermore, we demonstrate this has functional and physiological relevance as giantin KO zebrafish show ectopic calcificatied structures, similar to phenotypes seen in the human congenital disorder of glycosylation, HFTC.

We report that 24 enzymes involved in multiple glycosylation pathways exhibit altered expression following *GOLGB1* ablation. This implies that this change is not in response to a deficiency in a single reaction but a global adjustment of Golgi biochemistry. We consider this an adaptive response to giantin loss-of-function and it suggests a plasticity within the system that could have relevance to many processes including cell differentiation, tissue morphogenesis and responses to the extracellular environment. This is supported by the fact that KO cells and zebrafish are both viable and relatively unaffected by the transcriptional changes seen here. Indeed, lectin binding is largely equivalent in WT and KO cells suggesting that the new enzymatic equilibrium is broadly effective and the fidelity of glycosylation is largely maintained.

Genetic adaptation is an increasingly reported response to CRISPR-Cas9 generated mutations (Cerikan et al., 2016; Rossi et al., 2015). Such changes mask the original gene function but arguably better reflect disease states. Giantin depletion by siRNA has been reported to cause the specific redistribution of glucosaminyl (N-acetyl) transferase 3 (GCNT3) from the Golgi to the ER (Petrosyan et al., 2012). Expression of this gene was unaffected in our study but it is possible that perturbed transport of other enzymes instigated the transcriptional changes seen here. We found that giantin is not responsible for trafficking of GALNT3 to the Golgi, so transcriptional down-regulation of this enzyme at least is not the result of an anterograde trafficking defect. In addition, we do not detect any defects in cilia formation or function in giantin KO cells (Stevenson et al., 2017) or KO zebrafish (Bergen et al., 2017) despite robust phenotypes following acute knockdown (using RNAi (Asante et al., 2013; Bergen et al., 2017). Our previous work showed that knocking down giantin expression using RNAi resulted in defects in cilia formation and function (Asante et al., 2013; Bergen et al., 2017). To our surprise, giantin KO RPE-1 cells show no gross defects in ciliogenesis (Stevenson et al., 2017). Our RNAseq data and other experiments strongly suggest that RCAN2 compensates for loss of giantin in cilia length control (Stevenson et al., 2017).

Glycosylation is a seemingly robust process, with multiple compensatory mechanisms having been reported in response to gene loss. For example, loss of MGAT in T cells leads to the redistribution of sugar donors within Golgi cisternae to permit the synthesis of structurally dissimilar but bioequivalent glycans (Mkhikian et al., 2016). Interestingly, loss of MGAT expression does not result in major changes in the expression of other glycosyltransferases (Mkhikian et al., 2016). However, other work has shown that loss of one *N*-acetylglucosaminyltransferase can to lead to compensatory upregulation of a functionally equivalent isoform (Takamatsu et al., 2010). While these studies demonstrate the capacity of glycosylation for self-correction with respect to a single reaction, our data show for the first time the role of a non-enzymatic Golgi protein in global control of glycosylation.

The GALNT family of enzymes comprises extensive overlapping substrate specificities and so is a prime candidate for compensation (Bennett et al., 2012; Schjoldager et al., 2015). Indeed, five GALNTs are differentially expressed between WT and KO cells; GALNT1, GALNT3, GALNT12 and GALNT16 were down-regulated whilst GALNT5 was upregulated. Furthermore, staining with HPA lectin, which binds to the Tn antigen generated by GALNTs, was equivalent in WT and KO cell lines suggesting that the efficiency of this reaction was broadly maintained following these changes. Increased GALNT5 activity may therefore be sufficient to counter the loss of the other four enzymes, or the remaining GALNTs act collectively to ensure efficient O-glycosylation. The manifestation of HFTC-like phenotypes in giantin KO fish however is consistent with the idea that loss of GALNT3 cannot be fully compensated for over time with respect to specific substrates. This contrasts with other work showing that deletion of either GALNT1 or GALNT2, or ectopic expression of GALNT3, does not result in substantial changes in expression of the other GALNTs (Schjoldager et al., 2015). Glycosylation deficiencies for specific proteins have also been reported in prostate cancer cells where giantin is non-functional (Petrosyan et al., 2014).

The observed changes in expression of genes encoding Golgi-resident enzymes following loss of giantin expression suggests the existence of a Golgi-based quality control pathway for glycosylation. One interpretation of our data is that giantin itself is actively monitoring glycan synthesis or cargo transit and adjusting gene expression accordingly. Such organelle-based signalling circuits are not without precedent; the nutrient sensor mTORC1 can interact with and phosphorylate transcription factor EB (TFEB) on the surface of lysosomes during starvation to promote its nuclear translocation (Settembre et al., 2012). Giantin itself lacks enzymatic activity but it could function as a signalling platform in this context. MAPK, PKD and PKA signalling have all been shown to regulate Golgi activity (Farhan and Rabouille, 2011) but whether any of these pathways intersect with giantin and transcription remains to be determined. No obvious trafficking defects were detected in the KO cells at steady state, consistent with a function independent of vesicle tethering. This agrees with a report showing that, unlike known tethers, mitochondrial relocation of giantin does not result in vesicle tethering to the mitochondrial membrane (Wong and Munro, 2014). Nonetheless the possibility remains that the control of intra-Golgi traffic by giantin acts to ensure the accurate distribution of enzymes across the stack and this intersects with a signalling loop that directs expression of glycosyltransferases.

The lack of major structural changes in the Golgi apparatus in our KO cells is consistent with mouse KO models (Lan et al., 2016) and knockdown systems (Asante et al., 2013; Koreishi et al., 2013). It has been reported that the introduction of giantin into *Drosophila* S2 cells promotes clustering of Golgi stacks into pseudo-ribbons, implying a role in lateral tethering (Koreishi et al., 2013). If this is the case, then the lack of Golgi fragmentation in our KO models indicates that other golgins might fulfil this function in vertebrate systems. One notable phenotype that we observed was the presence of circularised Golgi structures following nocodazole treatment in giantin KO cells. This is counter-intuitive to a role in lateral tethering since removal of an inter-cisternal tether should reduce, rather than encourage, interactions between cisternal rims. Considering giantin has a predicted reach of 450 nm into the cytosol it is possible that instead it blocks interaction between similar membranes. During ribbon assembly, it would then need to be excluded from the rims of the stacks that are coming together. Alternatively, it may play a structural role in maintaining flat cisternae through homo- or heterotypic interactions or regulate lipid composition. We only see these circular structures following disassembly, suggesting larger Golgi ribbon structures may be under other physical constraints that maintain its linear conformation. If giantin does have a role in maintaining cisternal structure, changes in protein localisation or lipid packing in its absence could play a role in controlling glycosyltransferase expression. Relocation of GM130 to larger Golgi elements in nocodazole-treated giantin KO cells is consistent with its accumulation on the ‘old Golgi’ and with at least a partial compensation of function following loss of giantin.

Quality control mechanisms, such as may be active here, are well documented in the ER but to our knowledge only one study has looked at this specifically in the Golgi (Oku et al., 2011). This report found ten Golgi-relevant genes were upregulated in response to Golgi stress by virtue of a seven nucleotide *cis*-acting element within their promoters termed the Golgi apparatus stress response element (GASE) (Oku et al., 2011). We failed to identify these genes in our RNAseq analysis, nor was there any enrichment for promoters containing the GASE motif in our hits. It is therefore unlikely this pathway is active in our KO cells, but perhaps similar mechanisms exist to detect changes in the proteoglycome and adjust transcription accordingly.

Considerable variation exists between giantin KO animal models (Katayama et al., 2011; Lan et al., 2016). All, however, exhibit defects that could be attributed to changes in glycosylation, which in turn affects ECM deposition (Stanley, 2016; Tran and Ten Hagen, 2013). Changes in this process due to altered glycosyltransferase expression could thus underlie the broad chondrogenesis and osteogenesis phenotypes seen in rodent KO animals, whilst the diversity seen with regards to phenotypes likely reflects model specific modes of adaptation. The latter will be determined by tissue specific expression patterns, different developmental pathways, or differing compensatory mechanisms to produce bioequivalent glycans between species. Unlike our zebrafish mutants, HFTC-like phenotypes have not been reported in rodent giantin KO models. As HFTC is a late onset disorder, rodent models may not be able to develop HFTC-like characteristics, since these animals die at birth, whilst adult KO zebrafish are viable.

Overall, our work identifies a previously uncharacterised mechanism through which the Golgi can regulate its own biochemistry to produce a functional proteoglycome. Understanding the ability of cells to adapt and modulate glycosylation pathways through long term changes in gene expression has implications for normal development and disease pathogenesis in diverse contexts including congenital disorders of glycosylation (Jaeken, 2010), the onset and progression of cancer (Pinho and Reis, 2015), and long term health in terms of tissue regeneration and repair.

## Author contributions

NLS designed and performed experiments, analysed data and wrote the paper, DJB designed and performed experiments, analysed data and helped write the paper, RS helped with the zebrafish experiments, KARB, EK, and EM performed and analysed microCT experiments. CLH helped to design experiments, interpret data and write the paper. DJS conceived and managed the project, contributed to data analysis, and helped to write the paper.

## Acknowledgements

We would like to thank Franck Perez and Gaelle Boncompain for sharing the RUSH system with us, the Earlham Institute for the RNAseq analysis, Emily Wyatt for her contribution to the project and Andrew Herman and the UoB flow cytometry facility for help with cell sorting. Thanks to Martin Lowe for sharing reagents and for helpful discussions. We also thank the MRC and Wolfson Foundation for establishing the Wolfson Bioimaging Facility, and confocal microscopy was supported by a BBSRC ALERT 13 capital grant (BB/L014181/1). We would like to thank Tom Davies for his help with CT scanning. The project was funded by the MRC (MR/K018019/1), Wellcome Trust (099848/Z/12/Z), and Arthritis Research UK (19476 and 21211).

## Materials and Methods

All reagents were purchased from Sigma-Aldrich unless stated otherwise.

### Cell culture

Human telomerase-immortalised retinal pigment epithelial cells (hTERT-RPE-1, ATCC) were grown in DMEM-F12 supplemented with 10% FCS (Life Technologies, Paisley, UK). Cell lines were not authenticated after purchase other than confirming absence of mycoplasma contamination.

Transfections were performed using Lipofectamine 2000™ according to the manufacturer’s instructions (Invitrogen, Carlsbad, CA). Flag-GALNT3 was obtained from ViGene Biosciences (Cat# CH897457, Rockville, MD) Str-Kdel/Man-SBP-EGFP was a gift from Franck Perez (Institut Curie, Paris, (Boncompain et al., 2012)). For drug treatments cells were incubated with 5 μM nocodazole (Santa Cruz, Heidelberg, Germany) or 5 μM brefeldin A diluted in growth medium at 37°C then washed 3x with growth medium for recovery.

### Zebrafish husbandry and mutant alleles

London AB zebrafish were used and maintained according to standard conditions (Westerfield, 2000) and staged accordingly (Kimmel et al., 1995). Ethical approval was obtained from the University of Bristol Ethical Review Committee using the Home Office Project License number 30/2863. The *golgb1*^*Q2948X*^ and *golgb1*^*X3078*^ alleles are described in (Bergen et al., 2017).

### Genome engineering

RPE-1 cells were transfected as above with 1μg each of paired gRNAs HSL0001186601 (ACCTGAGCACGGCCCACCAAGG) and HSR0001186603 (GTCGTTGACTTGCTGCAACAGG) (obtained from Sigma) targeting the *GOLGB1* gene plus 0.1 μg pSpCas9n(BB)-2A-GFP (Addgene plasmid #48140 PX461 (Ran et al., 2013)). After 48 hours GFP-positive cells were sorted into 96 well plates, seeding one cell per well to generate clones. To identify mutations, genomic DNA was prepared using a Purelink^®^ genomic DNA mini kit (Invitrogen, Carlsbad, CA) and the region targeted by the gRNAs amplified by PCR (primers: forward 5’-CTGGGTCTGGTTGTTGTTGGT-3’ reverse 5’-GGTGTCATGTTGGTGCTCAG-3’; reaction mix: Taq DNA polymerase with thermopol^®^ buffer, 10 mM dNTP mix, 10 μM each primer and 2 μl genomic DNA; reagents from NEB (M0267L, N0447L)). PCR products were cloned into the pGEM^®^ T Easy vector according to the manufacturer’s instructions and sequenced using predesigned primers against the T7 promoter (MWG Eurofins).

### Antibodies, labelling and microscopy

Antibodies used: mouse monoclonal anti-giantin (full length, Abcam, Cambridge, UK, ab37266), rabbit polyclonal anti-giantin (N-terminus, Covance, CA, PRB-114C), rabbit polyclonal anti-giantin (C-term, gift from Martin Lowe), mouse anti-GM130 and mouse GMAP210 (BD Biosciences, Oxford, UK, BD 610823 & BD 611712), sheep anti-TGN46 (Bio-Rad, Hertfordshire, UK, AHP500), sheep anti-GRASP65 (gift from Jon Lane), rabbit anti-Sec23a (homemade, polyclonal), mouse anti-ERGIC53 (monoclonal clone G1/93, Alexis Biochemicals, ALX-804-602-C100), rabbit anti-TFG (Novus Biologicals, Cambridge, UK, IMG5901A), Sec16A (KIAA0310, Bethyl Labs, Montgomery, TX, A300-648A), ER stress antibody sampler kit (Cell Signalling, Hertfordshire, UK, 9956), mouse anti-tubulin, rabbit anti-GALNT3 and rabbit polyclonal anti-FLAG (Sigma, Dorset, UK, T5168, HPA007613 & F7425), CASP (gift from Sean Munro), mouse anti-GAPDH (Abcam, Cambridge, UK, ab9484), and sheep anti-GALNT3 (R&D systems, Abingdon, UK, AF7174). Lectins used: HPA biotinylated lectin (Fisher Scientific, Loughborough, UK, L11271), Biotinylated lectin kit I (Vector laboratories, Peterborough UK, BK-1000). HABP (Merck, Hertfordshire, UK, 385911).

For antibody labelling, cells were grown on autoclaved coverslips (Menzel #1.5, Fisher Scientific, Loughborough, UK), rinsed with PBS and fixed in MeOH for 4 minutes at −20°C. Cells were then blocked in 3% BSA-PBS for 30 minutes and incubated with primary then secondary antibody for 1 hour each, washing in between. Nuclei were stained with DAPI [4,6-diamidino-2-phenylindole (Life Technologies, Paisley, UK, D1306)] for 3 minutes and coverslips mounted in Mowiol (MSD, Hertfordshire, UK) or Prolong Diamond antifade (Thermo Fisher, Paisley, UK). For lectin labelling, cells were washed in PBS and fixed in 3% PFA-PBS for 10 minutes at room temperature (for lectins) or 10 minutes on ice plus 10 minutes at room temperature (for HABP). Cells were permeabilised in 1% (lectins) or 0.1% (HABP) TX-100 in PBS and blocked as above. Biotinylated lectins were diluted to 4 μg/ml in block and incubated with cells for 40 minutes whilst HAPB was diluted to 5 μg/ml and incubated overnight at 4°C. Cells were washed with PBS, incubated with giantin antibody for 15 minutes, washed and labelled with streptavidin-A568 and anti-rabbit A488 (Fisher Scientific, Loughborough, UK, S11226). Cells were DAPI stained and mounted as above.

Fixed cells were imaged using an Olympus IX70 microscope with 60x 1.42 NA oil-immersion lens, Exfo 120 metal halide illumination with excitation, dichroic and emission filters (Semrock, Rochester, NY), and a Photometrics Coolsnap HQ2 CCD, controlled by Volocity 5.4.1 (Perkin Elmer, Seer Green, UK). Chromatic shifts in images were registration corrected using TetraSpek fluorescent beads (Thermo Fisher). Images were acquired as 0.2 μm z-stacks.

For RUSH assays, cells were seeded onto 35-mm glass-bottomed dishes (MatTek, Ashland, MA) or coverslips and transfected 24 hr prior to assay; at T0 cells were treated with 40 μM biotin then imaged every 15 seconds as a single plane for up to 1 hr or fixed at specific time points and stained as above. Live widefield microscopy proceeded using an Olympus IX81 microscope with 60x 1.42 numerical aperture oil-immersion lens, Sutter DG4 illumination with excitation filters, and multi-pass dichroic and multi-pass emission filters (Semrock). Images were collected using an Orca Flash 2.8 sCMOS controlled using Volocity 5.4.2 (PerkinElmer). Cells were kept at 37°C for the duration of the imaging.

Quantification of Golgi structure from widefield images was performed using ImageJ software. Maximum projection images (GRASP65 channel) were generated from 0.2 μm z-stacks and thresholded before applying the “analyse particles” feature excluding objects <0.5 μm^2^ or on the edge of the field of view. Golgi cisternal length and curvature measurements taken from micrographs were again made with ImageJ using the segmented line and angle tools. Cisternae number and RUSH experiments were quantified manually and blind.

### EM

Cells were fixed in 2.5% glutaraldehyde, washed for 5 minutes in 0.1 M cacodylate buffer then postfixed in 1% OsO4/0.1 M cacodylate buffer for 30 minutes. Cells were washed 3x with water and stained with 3% uranyl acetate for 20 minutes. After another rinse with water, cells were dehydrated by sequential 10-minute incubations with 70, 80, 90, 96, 100 and 100% EtOH before embedding in Epon™ at 70°C for 48 hours. Thin 70 nm serial sections were cut and stained with 3% uranyl acetate then lead citrate, washing 3x with water after each. Once dried, sections were imaged using a FEI Tecnai12.

### Immunoblotting

Cells were lysed in RIPA buffer (50 mM Tris pH7.5, 300 mM NaCl, 2% Triton-X100, 1% deoxycholate, 0.1% SDS, 1 mM EDTA) and samples separated by SDS-PAGE followed by transfer to nitrocellulose membranes. Membranes were blocked in 5% milk-TBST or 3% BSA-TBST for antibody and lectin probes respectively. Primary antibodies/lectins diluted in block were incubated with membrane overnight and detected using HRP-conjugated secondary antibodies or streptavidin respectively (Jackson ImmunoResearch, West Grove, PA) and enhanced chemiluminescence (GE Healthcare, Cardiff, United Kingdom).

### Quantitative PCR

Total RNA was isolated from ventral bone and cartilage of juvenile *golgb1*^*Q2948X*^ and *golgb1*^*X3078*^ genotyped fish (60 and 63 dpf respectively, n=3 per genotype) using RNeasy mini kit (cat# 74104, Qiagen, Manchester, UK). Subsequently, a reverse transcriptase reaction was performed by using Superscript IV (cat# 18091050, Thermo Fisher). Zebrafish *galnt3* (XM_009300463.2) coding sequence was confirmed by multi-species nucleotide BLAST (NCBI) leading to *galnt3* forward 5’-TCCTTCAGAGTGTGGCAGTG and reverse 5’-TGATGGTGTTGTGGCCTTTA primers. *gapdh* as a reference gene was used (forward 5’-TGTTCCAGTACGACTCCACC and reverse 3’-GCCATACCAGTAAGCTTGCC). Quantitative Real-Time PCR (qPCR) reactions (quadruplicates per individual) using DyNAmo HS SYBR green (F410L, Thermo Fisher) with PCR cycles (40 times) of 95°C 25 seconds, 57.5°C 30 second, and 70°C 45 seconds followed by a standard melt curve were applied (QuantStudio3, Applied Biosystems).

### RNAseq

Triplicate samples of mRNA from giantin knockout cells and WT RPE-1 were analysed by RNAseq by the Earlham Institute (formerly The Genome Analysis Centre). The libraries were constructed by The Earlham Institute on a PerkinElmer Sciclone using the TruSeq RNA protocol v2 (Illumina 15026495 Rev.F). The library preparation involved the initial QC of the RNA using a Tecan plate reader with the Quant-iT™ RNA Assay Kit (Life technologies/Invitrogen Q-33140) and the Quant-iT™ DNA Assay Kit, high sensitivity (Life technologies/Invitrogen Q-33120). Finally, the quality of the RNA was established using the PerkinElmer GX with a high sensitivity chip and High Sensitivity DNA reagents (PerkinElmer 5067-4626). RNA quality scores were 8.7 and 9.8 for two of the samples and 10.0 (for the remaining 4 samples). 1 ug of RNA was purified to extract mRNA with a poly-A pull down using biotin beads, fragmented and the first strand cDNA was synthesised. This process reverse transcribes the cleaved RNA fragments primed with random hexamers into first strand cDNA using reverse transcriptase and random primers. The ends of the samples were repaired using the 3’ to 5’ exonuclease activity to remove the 3’ overhangs and the polymerase activity to fill in the 5’ overhangs creating blunt ends. A single ‘A’ nucleotide was added to the 3’ ends of the blunt fragments to prevent them from ligating to one another during the adapter ligation reaction. A corresponding single ‘T’ nucleotide on the 3’ end of the adapter provided a complementary overhang for ligating the adapter to the fragment. This strategy ensured a low rate of chimera formation. The ligation of a number indexing adapters to the ends of the DNA fragments prepared them for hybridisation onto a flow cell. The ligated products were subjected to a bead based size selection using Beckman Coulter XP beads (Beckman Coulter A63880) to remove un-ligated adapters, as well as any adapters that may have ligated to one another. Prior to hybridisation to the flow cell the samples were amplified by PCR to selectively enrich those DNA fragments that have adapter molecules on both ends and to amplify the amount of DNA in the library. The PCR was performed with a PCR primer cocktail that annealed to the ends of the adapter. The insert size of the libraries was verified by running an aliquot of the DNA library on a PerkinElmer GX using the High Sensitivity DNA chip and reagents (PerkinElmer CLS760672) and the concentration was determined by using the Tecan plate reader. The resulting libraries were then equimolar pooled and Q-PCR was performed on the pool prior to clustering.

These six total RNA samples were sequenced over two lanes and aligned against the human genome reference build 38 followed by differential expression analysis between the wildtype and knockout samples. QC was done using FastQC (version 0.11.2). An in-house contamination-screening pipeline (Kontaminant) was used to check for any obvious contamination in the raw reads. Since the data quality was good, there was no trimming done on the raw reads. Alignment of RNAseq reads to the human genome reference was done using TopHat (version 2.1.0) with “min-anchor-length” 12 and “max-multi hits” 20. The log_2_ of the fold-change was used in further analysis.

### Micro-Computed Tomography Scanning (μCT)

Female fish (n=3) carrying *golgb1*^*WT/WT*^ and mutant *golgb1*^*Q2948X/Q2948X*^ alleles were preserved in absolute ethanol at 8 mpf. Prior to scanning, the samples were packed in a polystyrene tube and scanned with a Bruker SkyScan 1272 (Kontich, Belgium) at a 21.8 or 4 μm resolution. The X-ray current was set at 200 μA with a voltage of 50 kV.

Eight zebrafish (four wild-type and four mutant *golgb1*^*X3078/X3078*^ fish) were scanned together using a Nikon X-TEK 225 HT computed tomography (CT) scanner at a resolution of 21 μm. Fish were arranged in a circle with seven on the outside surrounding the eighth fish. The first specimen was labelled so it was identifiable in the CT scans using a radiopaque sticker, and the remaining fish were numbered clockwise.

3D tomography images and movies for the golgb1^Q2948X^ allele were generated using CTvox software (v.3.0.0). For the golgb1^X3078^ allele and siblings the fish were segmented using Avizo (v. 9.3, Visualization Sciences Group), by thresholding greyscale values and labelling specific regions, bones and individual fish in materials for 3D visualisation. BMD was calculated relative to phantoms of known hydroxyl appetite density 0.25 and 0.75g/cm^3^ in 3 regions: for the skull, the lower jaw and vertebrae from multiple slices: 25 anteroposterior slices for the sagittal, the anterior-most 15 anteroposterior slices for the lower jaw, and the posterior half of the last vertebra before the end of the ribs. Using the Material Statistics module of Avizo, the mean greyscale value was calculated for each of these regions for each fish. BMD in each region was then calculated by multiplying the greyscale mean value by the maximum of 2.5 arbitrary units divided by the actual greyscale maximum of 65535. Finally, measurements of maximum lower jaw width and length were measured using the 3D CT reconstruction for both alleles.

### Quantification and statistical analysis

Statistical analyses were performed using GraphPad Prism 7.00. The tests used, n numbers and sample sizes are indicated in the figure legends, p-values are shown on the figures. All tests met standard assumptions and the variation between each group is shown. Sample sizes were chosen based on previous, similar experimental outcomes and based on standard assumptions. No samples were excluded. Randomisation and blinding were not used except where the genotype of zebrafish was determined after experimentation.

### Data availability

Raw RNAseq data are available in the ArrayExpress database (www.ebi.ac.uk/arrayexpress) under accession number E-MTAB-5618.

